# Multi-Omic Signatures of Sarcoidosis and Progression in Bronchoalveolar Lavage Cells

**DOI:** 10.1101/2023.01.26.525601

**Authors:** Iain R. Konigsberg, Nancy W. Lin, Shu-Yi Liao, Cuining Liu, Kristyn MacPhail, Margaret M. Mroz, Elizabeth Davidson, Clara I. Restrepo, Sunita Sharma, Li Li, Lisa A. Maier, Ivana V. Yang

**Author notes:** These authors contributed equally (Iain R. Konigsberg, Nancy W. Lin, Lisa A. Maier, Ivana V. Yang).

## Abstract

**Introduction:** Sarcoidosis is a heterogeneous, granulomatous disease that can prove difficult to diagnose, with no accurate biomarkers of disease progression. Therefore, we profiled and integrated the DNA methylome, mRNAs, and microRNAs to identify molecular changes associated with sarcoidosis and disease progression that might illuminate underlying mechanisms of disease and potential genomic biomarkers.

**Methods:** Bronchoalveolar lavage cells from 64 sarcoidosis subjects and 16 healthy controls were used. DNA methylation was profiled on Illumina HumanMethylationEPIC arrays, mRNA by RNA-sequencing, and miRNAs by small RNA-sequencing. Linear models were fit to test for effect of diagnosis and phenotype, adjusting for age, sex, and smoking. We built a supervised multi-omics model using a subset of features from each dataset.

**Results:** We identified 46,812 CpGs, 1,842 mRNAs, and 5 miRNAs associated with sarcoidosis versus controls and 1 mRNA, *SEPP1* - a protein that supplies selenium to cells, associated with disease progression. Our integrated model emphasized the prominence of the PI3K/AKT1 pathway in sarcoidosis, which is important in T cell and mTOR function. Novel immune related genes and miRNAs including *LYST, RGS14, SLFN12L*, and hsa-miR-199b-5p, distinguished sarcoidosis from controls. Our integrated model also demonstrated differential expression/methylation of *IL20RB, ABCC11, SFSWAP, AGBL4*, miR-146a-3p, and miR-378b between non-progressive and progressive sarcoidosis.

**Conclusions:** Leveraging the DNA methylome, transcriptome, and miRNA-sequencing in sarcoidosis BAL cells, we detected widespread molecular changes associated with disease, many which are involved in immune response. These molecules may serve as diagnostic/prognostic biomarkers and/or drug targets, although future testing will be required for confirmation.

## Introduction

Sarcoidosis is a heterogeneous disease characterized by non-caseating granulomatous inflammation that differentially impacts Black individuals and women^1^. The lungs are involved in over 90% of individuals^2^. Those with pulmonary sarcoidosis can be asymptomatic or demonstrate remission or resolution; however, progression can result in impairment and/or pulmonary fibrosis, the main cause of mortality^3^. The course of pulmonary sarcoidosis is unpredictable, with at least 25% of patients developing chronic or progressive disease requiring treatment^4^. Driven by a combination of genetic, environmental, and host immunologic factors, the underlying cause(s) of sarcoidosis are currently unknown. Multiple lines of evidence point towards an antigenic stimulus including HLA allele associations, environmental, seasonal, and regional patterns, and a Th1 predominate immune response in which CD4+ T cells secrete IFN-γ and TNF-α^5^. In addition, aberrant and dysfunctional immune responses are associated with sarcoidosis and supported by genome wide transcriptome studies^6^. While previous studies have elucidated many contributors to disease development and progression, many knowledge gaps remain.

Epigenetic mechanisms such as DNA methylation (DNAm) and microRNAs (miRNAs) mediate gene expression, are modified by exposures, and are dynamic and reversible, making them candidates for gene regulation in sarcoidosis as well as promising biomarkers and therapeutic targets. Epigenetic dysregulation has been identified in many lung diseases and likely drives sarcoidosis progression and manifestations. We have previously demonstrated changes in DNAm and mRNA gene expression in bronchoalveolar lavage (BAL) cells from chronic beryllium disease (a granulomatous lung disease caused by beryllium exposure) patients and a small sample of sarcoidosis patients^7,8^. No other studies have evaluated epigenome-wide DNAm in sarcoidosis although others have identified miRNAs associated with disease^9,10^. Additionally, BALF miRNAs such as miR-27b, miR-192, and miR-221 have been associated with pulmonary sarcoidosis progression^11^. These studies were limited by small sample sizes, targeted as opposed to genome-wide approaches, and a lack of integration of different omics modalities.

With a larger integrated approach utilizing several genomic data types, we hypothesize that epigenomic studies could link risk factors to disease pathobiology to better understand disease course, and subclassify patients based on molecular profiling; ultimately this would direct focused research on disease manifestations and treatment. In a first step, we conducted this study to profile genome-wide DNAm, mRNA and miRNA expression in sarcoidosis BAL cells, stratified by disease progression. Analyzing each dataset separately, we identified many molecular features associated with disease overall. By next constructing a sparse multi-omic model incorporating DNAm, mRNAs, and miRNAs, we identified features associated with sarcoidosis and pulmonary progression.

## Methods

### Study population

Sarcoidosis BAL cell samples were obtained from the Genomic Research in Alpha-1 Antitrypsin Deficiency and Sarcoidosis (GRADS) consortium^12^ (including National Jewish Health (NJH, n=17) and non-NJH GRADS cases (n=39)) and cases from the Granuloma Biorepository at NJH (n=8). Controls with no history of lung disease were obtained from the NJH Donor Lung Core (n=16). All sarcoidosis subjects met the ATS/ERS criteria^13^ for tissue biopsy confirmation of diagnosis of sarcoidosis. The non-progressive phenotype was defined as having either acute (i.e. consistent with Lofgren’s syndrome) or non-acute disease presentation, no new organ involvement, lung function testing with <10% decline in FVC or FEV_1_, <15% decline in DL_CO,_ and stable chest imaging within 2 years after BAL. The progressive phenotype had a non-acute disease presentation; lung function testing with ≥10% decline in FVC or FEV_1_; or ≥15% decline in DL_CO_; worsening chest imaging; and/or if they required initiation of systemic immunosuppressive treatment any time up to 2 years after BAL. Non-NJH GRADS cases were phenotyped based on disease acuity, PFT, and treatment status only. See Supplemental Methods for more details.

### Bronchoscopy and nucleic acid processing

Bronchoscopy with BAL was performed as previously described.^12,14^ Cells were isolated and frozen at −80C in RLT buffer. DNA and RNA were extracted using the Qiagen AllPrep DNA/RNA extraction mini kit. Purified genomic DNA was bisulfite-converted with the Zymo EZ-96 DNA Methylation bisulfite conversion kit, followed by whole-genome amplification and enzymatic fragmentation. DNA was denatured and hybridized to Illumina Infinium HumanMethylationEPIC BeadChips, followed by single base extension. Hybridized BeadChips were stained, washed, and scanned using Illumina’s iScan System. mRNA libraries were prepared from 500 ng total RNA with TruSeq stranded mRNA library preparation kits (Illumina) and miRNA libraries were constructed using Lexogen Small RNA-Seq library preparation kits. RNA libraries were sequenced at an average depth of 80M 150bp paired-end reads on the Illumina NovaSeq 6000. RNA-Seq reads from an additional 28 samples were obtained from the GRADS consortium^6^. Additional details are outlined in the supplemental methods.

### Data analysis

For each dataset, linear models were fit to each feature testing for an effect of sarcoidosis diagnosis or progressive vs. non-progressive disease while adjusting for age, sex, and smoking status. RNA-seq data was additionally adjusted for a surrogate variable capturing a batch effect of sequencing center^15^ (**Supplementary Figure 1**). Null test statistic distributions were derived for each analysis using bacon to reduce inflation^16^. *P*-values were adjusted to a 5% false discovery rate (FDR) to account for multiple testing using the Benjamini-Hochberg procedure^17^. Enrichment of significant results in the Gene Ontology (GO) resource^18^ and the Kyoto Encyclopedia of Genes and Genomes (KEGG)^19^ was performed using GOmeth^20^ for DNAm data and clusterProfiler^21^ for mRNA data. CpGs were annotated to CpG islands, shelves, and shores, and gene elements using the annotatr R package^22^. Experimentally confirmed miRNA target genes were obtained from MirTarBase^23^. For each miRNA, we retained target genes that had at least 2 sources of evidence (reporter assay, Western blot, qPCR, Microarray, NGS, pSILAC, CLIP-Seq, or other), including at least one source considered strong evidence (reporter assay, Western blot, or qPCR).

### Data Integration Analysis for Biomarker discovery using Latent cOmponents (DIABLO)

Methylation M values and normalized mRNA and miRNA gene expression values were used as input for a DIABLO^24^ model implemented in the mixOmics R package^25^. Gene expression values were normalized to library size using DESeq2^26^ and transformed to account for heteroscedasticity with a variance stabilizing transformation (VST)^27^. A subset of features from each dataset for 2 model components were selected for model input using LASSO regression and 5-fold cross validation repeated 10 times.

## Results

### Demographics of study population

We recruited 64 sarcoidosis cases, including 26 progressive and 38 non-progressive, and 16 healthy controls. Demographic information at time of BAL is displayed in **Table 1**. No significant differences were observed in race, ethnicity, or age. Non-progressive cases were more likely female than progressive cases. Progressive cases presented with significantly reduced FEV_1_ and FVC and more Stage 2 disease.

**Table 1.** Demographic information of control, progressive sarcoidosis, and non-progressive sarcoidosis cases at time of bronchoscopy with lavage.

|  | Control | Non-Progressive | Progressive | *p* |
| --- | --- | --- | --- | --- |
| n | 16 | 38 | 26 |  |
| Age (mean (SD)) | 55.62<br>(9.74) | 51.68<br>(9.88) | 51.19<br>(9.61) | 0.316* |
| Sex = Male (%) | 11 (68.8) | 11 (28.9) | 16 (61.5) | 0.006** |
| Race (%) |  |  |  | 0.242** |
| Asian | 1 (6.2) | 1 (2.6) | 0 (0.0) |  |
| Black | 0 (0.0) | 6 (15.8) | 5 (19.2) |  |
| White | 15 (93.8) | 31 (81.6) | 21 (80.8) |  |
| Ethnicity = Non-Hispanic (%) | 15 (93.8) | 36 (94.7) | 25 (100.0) | 0.441** |
| Smoking Status = Former (%) | 0 | 12 (31.6) | 6 (23.1) | 0.0275** |
| FVC (mean (SD)) |  | 97.42<br>(11.25) | 85.92<br>(14.21) | 0.001<sup>#</sup> |
| FEV<sub>1</sub> (mean (SD)) |  | 101.03<br>(12.19) | 86.40<br>(15.04) | <0.001* |
| DLCO (mean (SD)) |  | 87.71<br>(21.81) | 84.42<br>(12.64) | 0.491* |
| Scadding Stage (%) |  |  |  | <0.001** |
| 0 |  | 13 (34.2) | 1 (3.8) |  |
| 1 |  | 20 (52.6) | 1 (3.8) |  |
| 2 |  | 4 (10.5) | 22 (84.6) |  |
| 3 |  | 1 (2.6) | 2 (7.7) |  |

\*One-way ANOVA, \*\*Fisher's exact test, <sup>#</sup>Mann-Whitney U test. FVC: forced vital capacity. FEV<sub>1</sub>: forced expiratory volume over 1 second. DLCO: diffusing capacity of the lung for carbon monoxide.

### Gene expression representing immune pathways differs between cases and controls and to a lesser degree by disease progression phenotypes

We analyzed gene expression data to assess differences between both cases and controls and progressive and non-progressive disease. In the sarcoidosis-control comparison, we detected 1,842 differentially expressed genes (DEGs), of which 379 (20.6%) were significantly increased in sarcoidosis (**Figure 1A**; **Supplementary Table 1**). Top significant upregulated genes involved many genes relating to immunity, cell migration, and cell adhesion including: *EVL* (Enah/Vasp-like) and *SERPINA9* (serpin family A member 9). Most significant downregulated DEGs include *SFTPA2* and *SFTPC*, surfactant genes and transcription factors such as *CEBPD* (CCAAT Enhancer Binding Protein Delta) and *NKX2-1* (NK2 homeobox 1). A single DEG was found in progressive vs non-progressive cases: *SEPP1* (**Figure 1B**).

**Figure 1.**
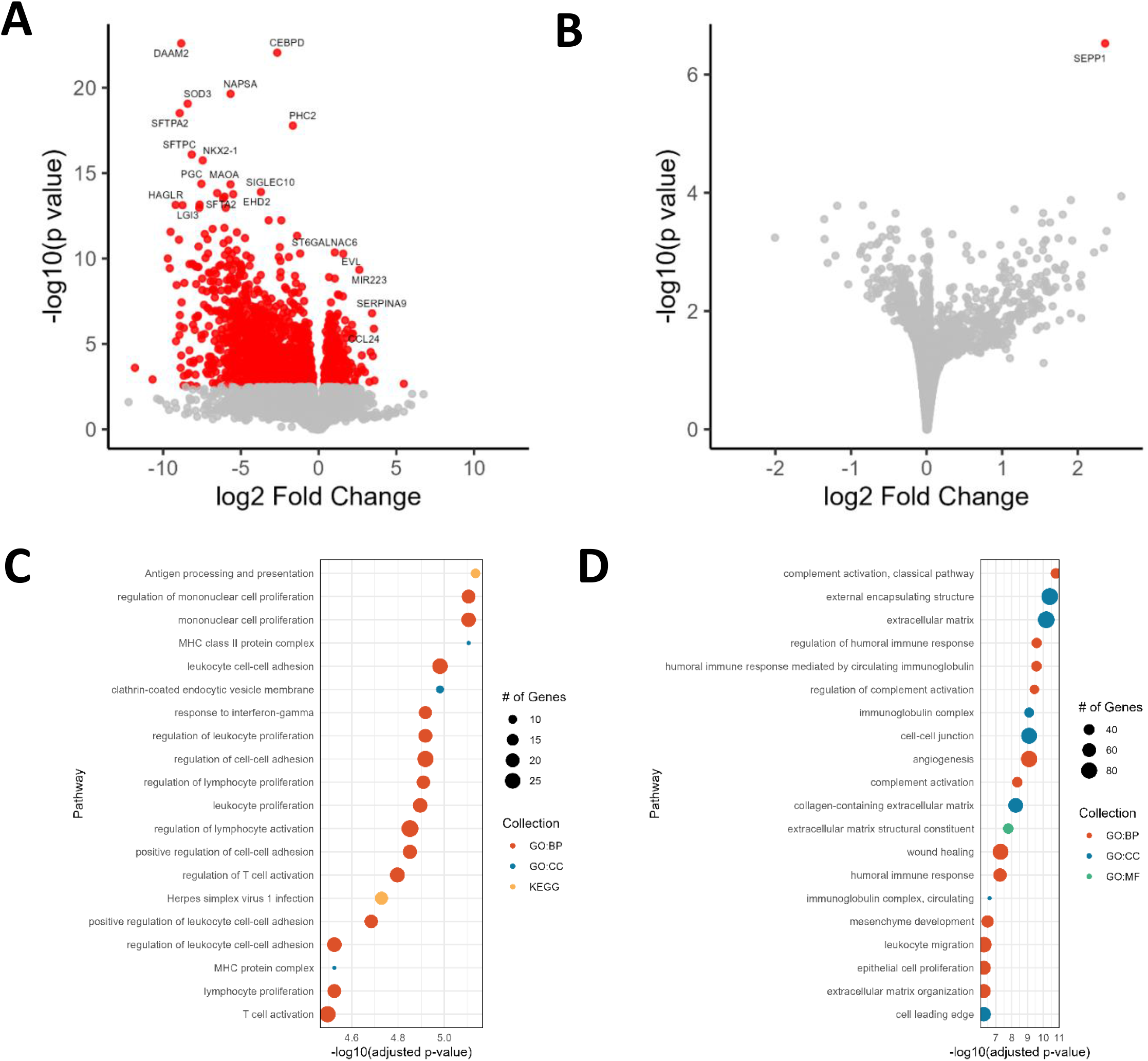
Differentially expressed genes in sarcoidosis. A) Differentially expressed genes in sarcoidosis vs. controls. B) Differentially expressed genes in progressive vs. non-progressive sarcoidosis. C) Pathway enrichment of upregulated mRNAs. D) Pathway enrichment of downregulated mRNAs.

We next tested whether sarcoidosis DEGs were overrepresented in GO^18^ and KEGG pathways using clusterProfiler^21^ (**Figure 1C-D**; **Supplementary Table 2**). Upregulated genes were enriched for 257 GO and 18 KEGG pathways. The most significantly enriched GO terms include MHC class II protein complex, regulation of mononuclear cell proliferation, leukocyte cell-cell adhesion, and response to interferon gamma. Top KEGG pathways include antigen processing and presentation, cell adhesion molecules, and multiple viral and autoimmune diseases including herpes simplex virus 1 infection, Epstein-Barr virus infection, viral myocarditis, Influenza A, tuberculosis, rheumatoid arthritis, and systemic lupus erythematosus. Downregulated genes were enriched for 512 GO and 0 KEGG pathways. The most significantly enriched GO terms include complement activation, classical pathway, extracellular matrix, regulation of humoral immune response, immunoglobulin complex, cell-cell junction, wound healing, mesenchyme development, and epithelial cell proliferation.

### DNA methylation differs between cases and controls and overlaps with gene expression

We next tested for differential methylation (DM) and identified 46,812 DNAm sites associated with sarcoidosis, of which 38,504 map to 15,208 unique genes (**Figure 2A**; **Supplementary Table 3**). We did not detect any DNAm sites significantly associated with disease progression. The majority of CpGs were hypermethylated in sarcoidosis (35,154; 75.1%). Significant probes were enriched for intronic regions (68.5% vs 60.8%; Fisher’s exact test *p* < 2.2 × 10^−16^) and FANTOM5 enhancers (9.17% vs 4.14%; Fisher’s exact test *p* < 2.2 × 10^−16^) relative to all tested DNAm sites. Hypomethylated DM sites were significantly enriched for 687 GO terms, with top hits including immune response, leukocyte activation, and defense response, and 88 KEGG pathways including viral protein interaction with cytokine and cytokine receptor, chemokine signaling pathway, Th17 cell differentiation, and T cell receptor signaling pathway (**Figure 2B**; **Supplementary Table 4**). Hypermethylated DM sites were significantly enriched for 146 GO terms, with top hits including actin cytoskeleton organization and 29 KEGG pathways including Rap1 signaling, Ras signaling, and phosphatidylinositol signaling system (**Figure 2C**). The most significant hypomethylated DM sites include CpGs that map to a predicted promoter region of *RGS14*, an intronic enhancer of *FGF18*, and *MGAT1*. Top hypermethylated DM sites mapped to genes such as *LYST* (lysosomal trafficking regulator), which regulates protein trafficking to lysosomes.

**Figure 2.**
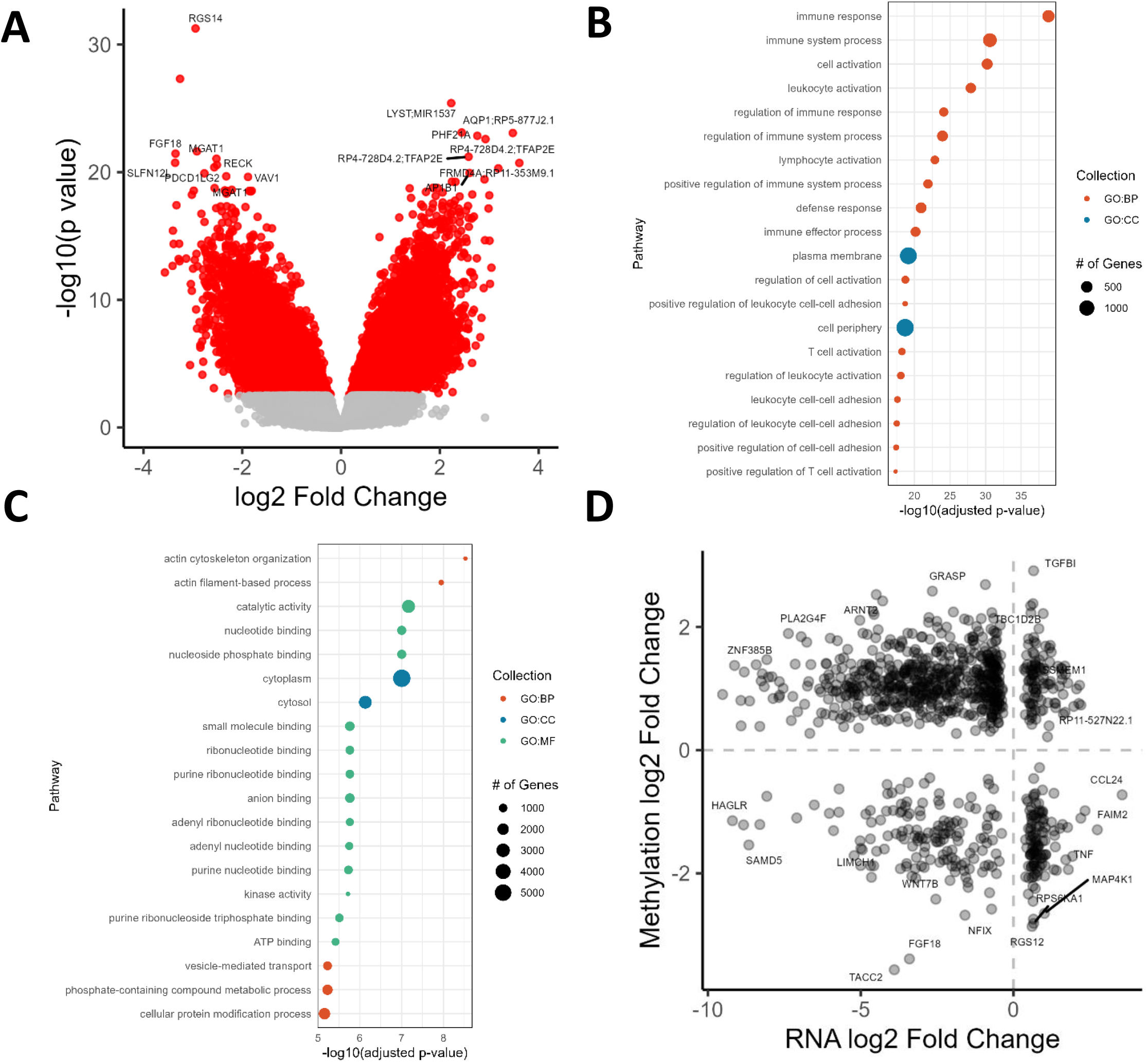
Differentially methylated sites in sarcoidosis. A) Differentially methylated sites in sarcoidosis vs. controls. B) Pathway enrichment of hypomethylated DNAm sites. C) Pathway enrichment of hypermethylated DNAm sites. D) Differentially expressed genes with DM DNAm sites.

We next overlapped DEGs and differentially methylated probes (DMPs) in sarcoidosis vs controls based on gene ID. 961 DEGs (52.2%) showed DM of 12,297 unique CpGs. 52.9% of associations (41,674/78,761) showed canonical inverse relationships: the directionality of DNAm was opposite that of gene expression (**Figure 2D**). Genes with canonical relationships and upregulated RNA include *TGFB1*. Canonical relationships with downregulated RNA include *AQP1, FRMD4A, ARNT2*, and *BMP4*. Genes with non-canonical relationships and downregulated RNA include *ITGA9*, an integrin component of receptor for *VCAM1, LIMCH1*, which positively regulates stress fiber assembly and stabilizes focal adhesions, and *FGF18*.

### miRNA expression differs between cases and controls and targets DEGs

We next compared miRNA expression in sarcoidosis vs controls and identified 5 miRNAs (hsa-miR-143-3p, hsa-miR-199a-3p/hsa-miR-199b-3p, hsa-miR-199b-5p, hsa-miR-582-3p & hsa-miR-582-5p) downregulated in sarcoidosis (**Figure 3A-B**; **Supplementary Table 5**). No miRNAs were associated with progression. We derived experimentally validated target genes for each significant miRNA using MirTarBase^23^. We identified 67 target genes, of which 4 (*AKT1, CD44, JAG1, PTGS2*) are targeted by 2 DE miRNAs (**Figure 3C**), and a further 15 were DEGs in our mRNA analysis, including *TNF*.

**Figure 3.**
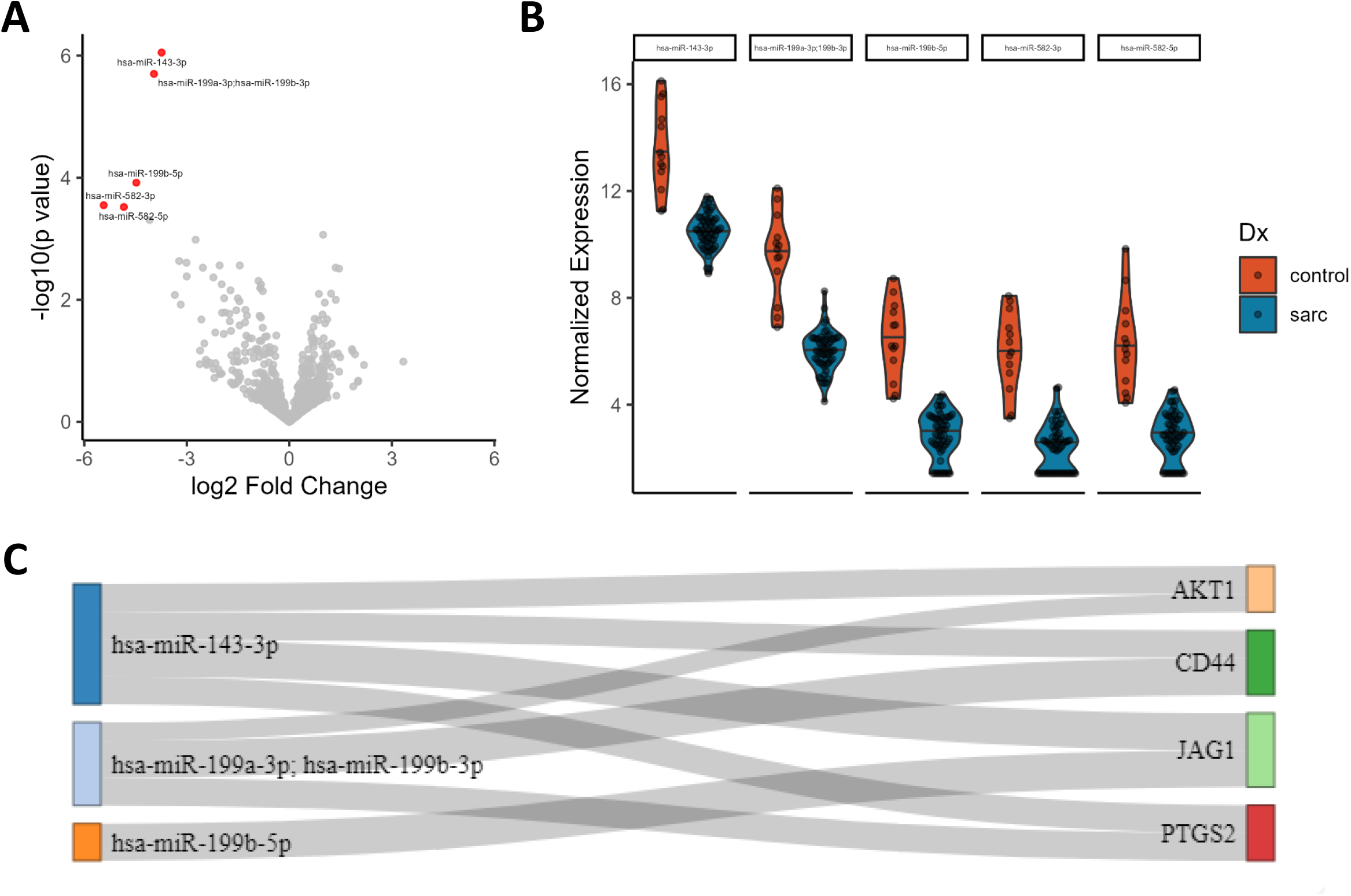
Differentially expressed microRNAs in sarcoidosis. A) Differentially expressed miRNAs in sarcoidosis vs. controls. B) Distribution of significant miRNAs’ expression in cases and controls. C) Sankey plot connecting DE miRNAs to target genes targeted by >1 DE miRNA. Connection width represents number of sources confirming relationship.

### Integrated model reveals associations with sarcoidosis and with progression not found in individual analyses

We used DIABLO to integrate the 3 datasets, including individuals present in all datasets after QC (n = 65; 13 controls, 19 progressive, 33 non-progressive sarcoidosis). With a goal to determine molecular features separating 3 groups (controls, progressive, and non-progressive sarcoidosis), we constructed a multi-omic model with 2 latent variables, which are linear combinations of input features. Using 5-fold cross validation repeated 10 times, we determined that selecting 5 DNAm sites, 2 mRNAs, and 13 miRNAs for latent variable 1 and 2 DNAm sites, 2 mRNAs, and 14 miRNAs for latent variable 2 maximized the prediction accuracy of the model and resulted in clustering based on diagnosis (**Figure 4A**; **Supplementary Table 6**). Latent variable 1 separates controls from sarcoidosis samples and latent variable 2 separates progressive from non-progressive sarcoidosis. Features included in each latent variable are shown in **Figure 4B** and **Supplementary Table 7**. We further constructed networks based on correlations of feature weights for each latent variable (**Figure 4C**). The two mRNAs contributing to latent variable 1 were *SFTPB* and *SFTPD*. DNAm sites contributing to latent variable 1 include DNAm sites hypermethylated in cases cg16962115, within an intron of *LYST*, and cg05300241 within an intron of *PIK3CD*. The remaining 3 DNAm sites were hypomethylated and included cg21949194 within an enhancer of *SOS1* (SOS Ras/Rac guanine nucleotide exchange factor 1), cg11370586 within the predicted promoter of *RGS14*, and cg03526142 within an exon of *SLFN12L*. All DNAm sites contributing to latent variable 1 were DMPs in our previous methylation analysis. In addition to the DE miR hsa-miR-199b-5p, miRNA features on latent variable 1 include hsa-miR-204-5p. Features contributing to latent variable 2 include the DNAm sites cg05479174, within an exon of *SFSWAP*, and cg06635176 within an intron of *AGBL4. IL20RB* (interleukin 20 receptor subunit beta) mRNA was upregulated in non-progressive sarcoidosis, while ABCC11 (ATP binding cassette subfamily C member 11) expression was higher in progressive sarcoidosis. miRNAs contributing to latent variable 1 include hsa-miR-146a-3p and hsa-miR-378b.

**Figure 4.**
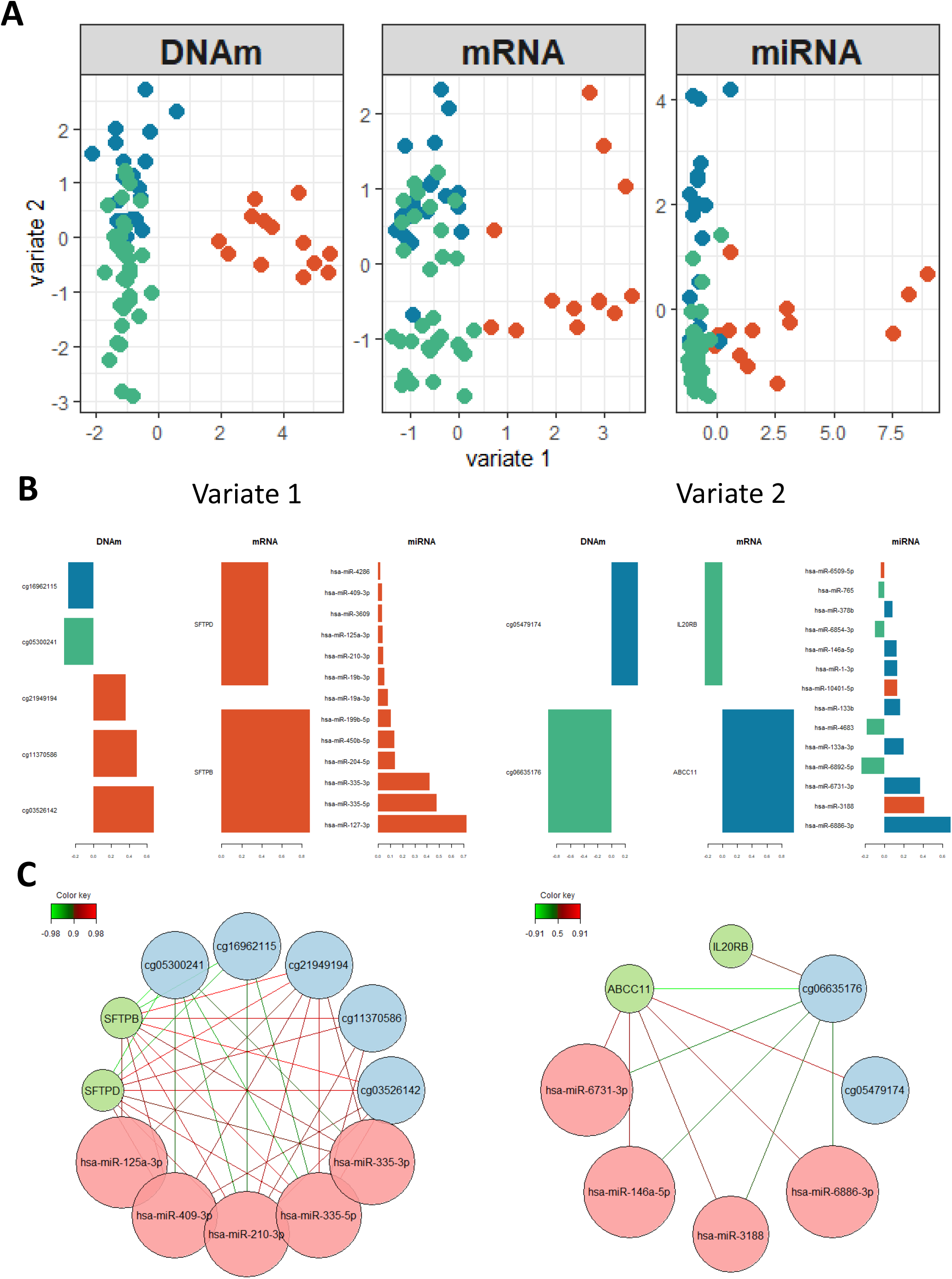
A sparse multi-omic model of sarcoidosis and progression. A) Projection of samples based on selected feature weights. B) Weights of selected features for latent variables. C) Network constructed from feature correlations on latent variables.

## Discussion

In this study, we present the first application of multiomic integration in sarcoidosis, leveraging coding and miRNA expression with DNA methylation data to construct a multiomic network. Through initial genome-wide profiling of DNA methylation, mRNA expression, and miRNA expression in sarcoidosis BAL, we identified 1,846 DEGs, 46,812 DMPs, and 5 DE miRNAs in sarcoidosis relative to healthy controls, as well as 1 DEG associated with sarcoidosis progression. By integrating omic datasets, we define pathogenic molecules/genes in sarcoidosis not identified using conventional modeling methods in single-omics datasets. While our single-omics approach only demonstrated 1 DEG for progression, the multiomic model identified several progression-associated methylation, mRNA, and miRNA features. We identify previously reported molecules and pathways associated with disease as well as implicate novel molecular features as potential drivers and modifiers of sarcoidosis, thus demonstrating the potential of integrative approaches.

Our single-omic analyses implicate multiple genes in the pathogenesis of sarcoidosis, with many of these genes involved in processes relating to inflammation and immunity (potentially associated with those involved in viral/atypical bacterial infections and autoimmunity) and extracellular matrix signaling. While we detected many associations with case status, we detected only a single omic association with progression: *SEPP1* mRNA levels. Results from the GRADS study demonstrated that *SEPP1* levels were inversely correlated with DLCO (%) and FVC (% pred) in sarcoidosis individuals^6^. These finding support a progressive phenotype, including lung function changes that we and others have used in the definition of progressive pulmonary disease.

Our study also demonstrated 5 DE miRNAs downregulated in sarcoidosis cases in the first study of genome-wide miRNAs in sarcoidosis. While these associations are novel, multiple target genes of these miRNAs have been implicated in sarcoidosis previously, including the genes *AKT1, CD44, JAG1*, and *PTGS2*. For example, CD44 has been found in areas of granuloma formation and fibrosis^28^ and is differentially expressed between Lofgren syndrome versus non-Lofgren syndrome subjects^29^. Through integrating genes with evidence of both DE and DM, we identified biologically relevant genes in sarcoidosis versus controls, including *HIF1A* and *TGFB1*. A previous study showed that HIF1A protein and mRNA levels were decreased in sarcoidosis granulomas^30^. We and others have found alterations in *TGFB1*, which encodes TGF-β, in pulmonary sarcoidosis^31,32^. Additionally, TGF-β genotypes have been associated with sarcoidosis severity^31^.

Our integrated model revealed several novel genes and miRNAs between sarcoidosis and controls, including *LYST, RGS14, SLFN12L*, and hsa-miR-199b-5p. The first 3 genes appear to have important roles in immune function. Specifically, *LYST*, a regulator of endosome/lysosome trafficking, can regulate TLR3 and TLR4 mediated pathways^33^, genes involved in the innate immune system which have been implicated in sarcoidosis^6,34,35^. *RGS14* is expressed in lymphocytes and regulates chemokine receptors to control immune responses to exogenous agents^36^. Finally, *SLFN21L* regulates thymocyte development and is downregulated in T-cell activation, suggesting a role as an immune response regulator^37^.

Our integrated analyses demonstrated miRNAs DE between progressive/non-progressive sarcoidosis including those previously described. Specifically, miR-146a-3p upregulated in progressive sarcoidosis is an indicator of inflammation and oxidative stress that may target *TLR4* and was previously found elevated in sarcoidosis BALF, as well as serum^38,39^. miR-378b, which was upregulated in progressive cases, was previously found associated with sarcoidosis^40^. We also identified novel genes including *IL20RB, ABCC11, SFSWAP* and *AGBL4*. Interestingly, *IL20RB* expression was increased in non-progressive versus progressive sarcoidosis in our study, suggesting that the activation of this inflammatory pathway is increased in the non-progressive phenotype. *ABCC11* is a gene that influences macrophage differentiation and induces TNF-α and IL17 through TLR4 signaling^41^; these results as well as the novel findings above support the importance of innate immune response genes in sarcoidosis.

Both our individual and integrative analyses identify molecules involved in PI3K/AK1 signaling, a pathway already recognized in sarcoidosis pathogenesis. For example, we observe hypermethylation of *PI3KCD* (phosphatidylinositol-4,5-bisphosphate 3-kinase catalytic subunit delta), which encodes a component of PI3K, and complexes with AKT1 to impact T cell differentiation and function^42^. Our miRNA results demonstrate *AKT1* as a target of downregulated miRNAs (miR-143-3p and miR-199a/miR-199b) in sarcoidosis. Both downregulation and upregulation of the PI3K/AKT signaling pathways have been associated with sarcoidosis^42,43^. Previous studies demonstrated both reduced proliferative response and exhaustion of T cells in progressive sarcoidosis is thought to be driven in part by inhibition of PI3K/AKT1 signaling^42^. The PI3K/AKT1 pathway has also been implicated in activation of mTOR^43^, which has been associated with granuloma formation^44^. Interestingly, in the recent GRADS study, PI3K activation was associated with an endotype of sarcoidosis characterized by hilar lymphadenopathy, pulmonary reticulation, less multiorgan involvement, and more environmental associations^6^.

While our sample size is larger than most previous sarcoidosis omic studies, power in our analyses was limited, especially in our phenotype analyses. In future studies, it will be important to both increase sample sizes and investigate associations with better-powered quantitative measures related to disease such as pulmonary function testing variables. We were unable to correct for cell proportions in sarcoidosis subjects relative to controls as our controls lacked cell differentials. Altered cell proportions may explain the downregulated epithelial genes in sarcoidosis BAL (e.g. decreased surfactant proteins in sarcoidosis) and some of the DNAm sites and DEGs detected in sarcoidosis versus control comparisons, although in general epithelial cells are not a large proportion of BAL. Finally, our multiomic models did not take into account demographic information. Regressing out demographic information such as age and sex may lead to more predictive networks. Despite these shortcomings, we detected widespread molecular changes and constructed multiomic networks associated with sarcoidosis as well as progression. Molecules discovered in these analyses shed light on disease pathogenesis and may also be leveraged therapeutically or as biomarkers after replication/validation in additional populations.

---

Supplementary Figure 1. Accounting for RNA-Sequencing batch effect. A) Principal component analysis of mRNA data, colored by sequencing location. B) Surrogate variable analysis of mRNA data, colored by sequencing location. C) Principal component analysis of mRNA data after regressing out SV1.

Supplementary Table 1. Differentially expressed genes associated with sarcoidosis.

Supplementary Table 2. Pathway over-representation analysis of DEGs.

Supplementary Table 3. Differentially methylated probes associated with sarcoidosis.

Supplementary Table 4. Pathway over-representation analysis of DMPs.

Supplementary Table 5. Differentially expressed miRNAs associated with sarcoidosis.

Supplementary Table 6. DIABLO cross-validation balanced error rates.

Supplementary Table 7. DIABLO loadings.

