## Supplementary Methods for "Multi-Omic Signatures of Sarcoidosis and Progression in Bronchoalveolar Lavage Cells"

Supplemental Methods

Study Population

For all NJH cases, medical records were reviewed for clinical features including acuity of presentation (acute/non-acute), organ involvement, pulmonary function tests (PFT), chest imaging and immunosuppressive treatment at time of BAL and up to two years after BAL. For the non-NJH GRADS cases, acuity of presentation, PFTs, and immunosuppressive treatment were available at and up to 6 months after BAL. Cases with fibrotic/Scadding stage 4 chest radiographic disease and/or on immunosuppressive treatment at time of BAL were excluded. Sarcoidosis cases were categorized as non-progressive or progressive pulmonary phenotypes.

Data processing & QC

Raw RNA-Seq files were obtained from GRADS. RNA paired-end reads from both GRADS and NJH samples were aligned at the gene level to Ensembl GrCh38 using STAR^1^. The forward read of miRNA FASTQ files were aligned to sequences from miRBase v22.1^2-7^ using miR-MaGiC^8^. miRNAs present in at least 50% of samples were retained (n = 802). Two samples were excluded from analysis as they were >3 standard deviations from the mean of either of the top 2 principal components of the data. Illumina idat signal intensity files were processed using SeSAMe; within-sample normalization with out-of-band probes and dye bias correction were performed^9^. Probes with non-unique mapping and off-target hybridization were removed. Additionally, probes with an average detection *p-*value ≥0.05 within samples and sex chromosome probes were removed prior to analysis. Methylation levels were analyzed as M values, and are presented in results as Beta values^10^.

1.	Dobin, A., Davis, C.A., Schlesinger, F., Drenkow, J., Zaleski, C., Jha, S., Batut, P., Chaisson, M., and Gingeras, T.R. (2013). STAR: ultrafast universal RNA-seq aligner. Bioinformatics *29*, 15-21. 10.1093/bioinformatics/bts635.

2.	Kozomara, A., Birgaoanu, M., and Griffiths-Jones, S. (2019). miRBase: from microRNA sequences to function. Nucleic Acids Res *47*, D155-D162. 10.1093/nar/gky1141.

9.	Zhou, W., Triche, T.J., Jr., Laird, P.W., and Shen, H. (2018). SeSAMe: reducing artifactual detection of DNA methylation by Infinium BeadChips in genomic deletions. Nucleic Acids Res *46*, e123. 10.1093/nar/gky691.

10.	Du, P., Zhang, X., Huang, C.-C., Jafari, N., Kibbe, W.A., Hou, L., and Lin, S.M. (2010). Comparison of Beta-value and M-value methods for quantifying methylation levels by microarray analysis. BMC Bioinformatics *11*. 10.1186/1471-2105-11-587.
