## Supplementary figures and images for "Multi-Omic Signatures of Sarcoidosis and Progression in Bronchoalveolar Lavage Cells"

### Supplementary Figure 1

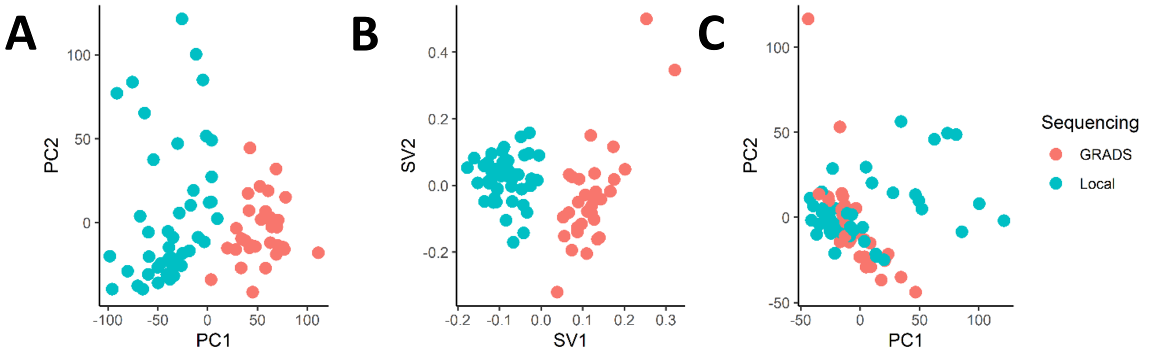
